# A coarse-grained DNA model to study protein-DNA interactions and liquid-liquid phase separation

**DOI:** 10.1101/2023.05.19.541513

**Authors:** Utkarsh Kapoor, Young C. Kim, Jeetain Mittal

## Abstract

Recent advances in coarse-grained (CG) computational models for DNA have enabled molecular-level insights into the behavior of DNA in complex multiscale systems. However, most existing CG DNA models are not compatible with CG protein models, limiting their applications for emerging topics such as protein-nucleic acid assemblies. Here, we present a new computationally efficient CG DNA model. We first use experimental data to establish the model’s ability to predict various aspects of DNA behavior, including melting thermodynamics and relevant local structural properties such as the major and minor grooves. We then employ an all-atom hydropathy scale to define non-bonded interactions between protein and DNA sites, to make our DNA model compatible with an existing CG protein model (HPS-Urry), that is extensively used to study protein phase separation, and show that our new model reasonably reproduces the experimental binding affinity for a prototypical protein-DNA system. To further demonstrate the capabilities of this new model, we simulate a full nucleosome with and without histone tails, on a microsecond timescale, generating conformational ensembles and provide molecular insights into the role of histone tails in influencing the liquid-liquid phase separation (LLPS) of HP1α proteins. We find that histone tails interact favorably with DNA, influencing the conformational ensemble of the DNA and antagonizing the contacts between HP1α and DNA, thus affecting the ability of DNA to promote LLPS of HP1α. These findings shed light on the complex molecular framework that fine-tunes the phase transition properties of heterochromatin proteins and contributes to heterochromatin regulation and function. Overall, the CG DNA model presented here is suitable to facilitate micron-scale studies with sub-nm resolution in many biological and engineering applications and can be used to investigate protein-DNA complexes, such as nucleosomes, or LLPS of proteins with DNA, enabling a mechanistic understanding of how molecular information may be propagated at the genome level.

## 1. Introduction

DNA is ubiquitous in biological systems. The assembly, compaction, and proper packaging of eukaryotic DNA into chromatin are critical components of cellular functions such as transcription and replication. A wide range of human diseases has been associated with defects in chromatin structure. Thus, understanding the molecular factors that govern chromatin organization is central to molecular biology, biophysics, and ultimately human health (1–4). Recently, liquid-liquid phase separation (LLPS) has been proposed as a mechanism for chromatin organization. Nucleosomes, which represent the basic subunits of chromatin structure, have been shown to localize in liquid-like droplets formed by Heterochromatin protein 1 (HP1)α protein (5–7).

In the context of proteins, simplified computational models have provided valuable mechanistic insights on driving forces underlying protein LLPS and have aided in elucidating molecular interactions in the condensed phases of proteins.(8) On the other hand, in the context of DNA, several coarse-grained (CG) models have been proposed: low-resolution CG models that can recapitulate the thermodynamics of DNA hybridization but lack the information needed to capture the structural details (9–14), or higher resolution CG models that can simultaneously reproduce thermodynamic, mechanical, and structural properties of DNA and RNA reasonably well (15–28).

In studies employing high-resolution CG models for both DNA and proteins with amino-acid level detail, the CG force field relies on protein-DNA cross-interaction parameters that are often non-transferable and system-specific. These parameters are typically re-parameterized to reproduce experimental measurements, such as the molar dissociation constant, *K_d_* (28–32). The use of complex potential energy functions, which include anisotropic potentials between bases involved in base pair stacking and hydrogen bonding interactions, makes these computationally expensive, even with the latest state-of-the-art computer software and hardware. In the context of chromatin, previous studies examining nucleosome structure and dynamics have either been limited in length- and time-scale (33–36) or have ignored critical molecular features of nucleotides and their specific interactions with different amino acids (37–47).

In this paper, we present a CG DNA model designed for the study of protein-DNA interactions and their impact on protein LLPS. Our CG model provides a reasonable description of various aspects of double-stranded DNA (dsDNA), including Watson-Crick base pairing, melting, hybridization, and major and minor grooves. To demonstrate the suitability of our model to perform large-scale simulations, we conduct simulations of a mono-nucleosome over long timescales. This allows us to gain insights into the influence of histone tails on the conformational properties of DNA when bound to the histone core. Additionally, we utilize the model to investigate the co-localization of nucleosome within the condensed phase of HP1α.

The remainder of this article is organized as follows: First, we provide a detailed description of the CG DNA model. Next, we outline the protocols and simulation methods employed for the parametrization and validation of the model, along with the presentation of parametrization results. To evaluate the model’s accuracy, we investigate its ability to predict various experimentally observed phenomena, including DNA melting behavior and structural properties of dsDNA. Subsequently, we establish non-bonded protein-DNA interactions using an all-atom **h**ydro**p**athy **s**cale (HPS) and demonstrate the compatibility of the CG DNA model with the previously proposed CG HPS-Urry protein model (48–50). Finally, we explore a range of conformational states of disordered histone tails to gain insights into how the presence of histone tails may modulate the behavior of nucleosomal DNA. We also investigate the role histone tails in modulating the LLPS of HP1α proteins. In Conclusions, we summarize the key findings of our study and discuss potential future applications.

## 2. Model and Methods

### 2.1. Development of CG model for dsDNA

#### 2.1.1. Physical representation

Our 2-bead CG DNA model, shown in **Figure 1**, maps nucleotide to two interaction sites where one bead represents the sugar-phosphate backbone carrying an overall -1 charge, connected to neighboring backbone beads via harmonic bonds, and the other represents the base connected to the corresponding backbone bead via harmonic bonds. The model differentiates between bases ADE (A), THY (T), CYT (C), and GUA (G) in their stacking and hydrogen bonding interactions.

**Figure 1.**
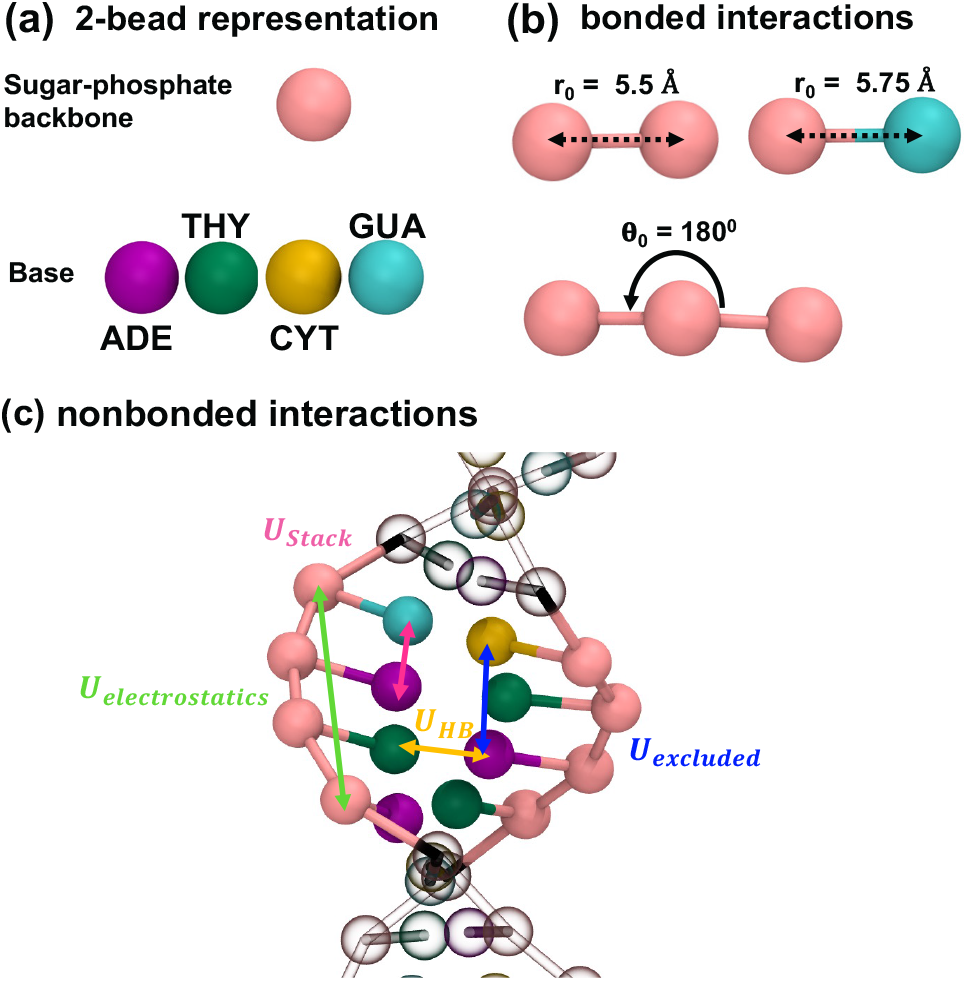
Schematic representation (not drawn to scale) of the 2-bead CG DNA model proposed in this work. The model includes a sugar-phosphate backbone bead (pink color), and bases A, T, C, G represented by green, violet, yellow, and cyan colors, respectively.

The potential energy of the system includes six distinct contributions:

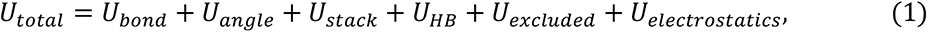

where *U_bond_* and *U_angle_* are the bonded potentials for intra-molecular bonds and angles, respectively, *U_stack_* is the stacking potential between base beads within the same strand, *U_HB_* is the hydrogen bonding (HB) potential between complementary base beads, *U_excluded_* is the excluded-volume potential, and *U_electroststics_* is the screened Coulomb potential between charged backbone beads (see **Figure 1c**). The details of each term are described in the following subsection. The mass of each bead (in amu) is taken from the corresponding the all-atom structure. Specifically, the masses of the sugar-phosphate backbone, as well as the A, T, C, and G base beads are 178.08 amu, 134.1 amu, 125.1 amu, 110.1 amu, and 150.1 amu, respectively.

#### 2.1.2. Bonded interactions

The first two terms in eq. 1 represent the standard intramolecular bond and angle potentials. The bonds between the backbone-backbone and backbone-base beads are described using a harmonic potential:

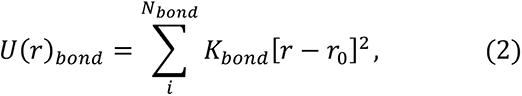

where *N_bond_* represents the number of bonds, *K_bond_* is the spring constant, and *r*_0_ is the equilibrium bond length. The three-body angle potential is applied solely to three consecutive backbone beads and is defined by a cosine/squared-style harmonic potential:

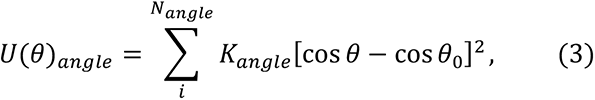

where *N_angle_* represents the number of angles, *K_angle_* is the spring constant, and *θ*_0_ denotes the equilibrium angle. The values of the equilibrium bond lengths, denoted as *r*_0_, for both backbone-backbone and backbone-base beads, along with the force constants *K_bond_* and *K_angle_* are summarized in **Table 1**. It is worth mentioning that the values of the equilibrium bond lengths are drawn from the canonical B-form of DNA, and the force constant *K_angle_* has been empirically optimized to maintain the structural properties of the dsDNA.

**Table 1:** Bonded interaction parameters associated with eqs. 2 and 3.

|  |  |  |  |
| --- | --- | --- | --- |
| **Bonds** | backbone-backbone | $K_{bond}$ | 50 kcal/mol/Å<sup>2</sup> |
| | | $r_0$ | 5.5 Å |
| | backbone-base | $K_{bond}$ | 50 kcal/mol/Å<sup>2</sup> |
| | | $r_0$ | 5.75 Å |
| **Angles** | backbone-backbone-backbone | $K_{angle}$ | 40 kcal/mol |
| | | $\theta_0$ | 180 degrees |

#### 2.1.3. Non-bonded interactions

The remaining terms in eq. 1 describe different pairwise non-bonded interactions. It should be noted that non-bonded interactions between directly bonded beads are excluded. The potentials *U_stack_*, *U_HB_* and *U_excluded_* are mutually exclusive, implying that a pair of beads contributes solely to one of these terms. The *U_stack_*, term captures the base stacking between consecutive base beads (i.e., beads *i* and *i*+1) within the same strand. The stacking potential is represented by a 12-10 Lennard-Jones (LJ) potential:

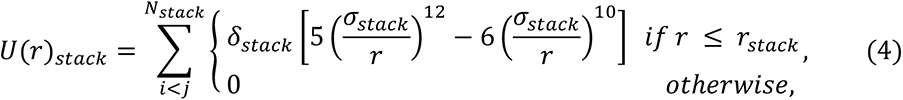

where *N*_stack_ represents the number of stacking pairs, *δ*_stack_ corresponds the stacking interaction strength, *σ*_stack_ denotes the equilibrium stacking distance, and *r*_stack_is the cutoff distance for the stacking potential. The *U*_*HB*_ term represents the interaction between complementary Watson-Crick base pairs. This potential is applied to both intra- and inter-strand base beads, except for the nearest and next-nearest intra-strand base beads (i.e., |*i* − *j*| ≤ 2, where *i* and *j* are the corresponding backbone bead indices). Similar to the stacking potential, the 12-10 LJ potential is used to model this interaction:

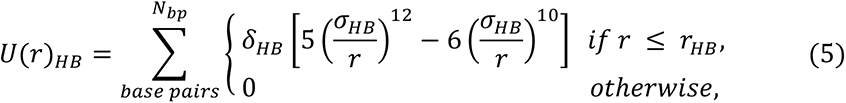

where *N*_*bp*_ represents the number of HB pairs, *δ*_*HB*_ corresponds to the HB interaction strength, *σ*_*HB*_ denotes the equilibrium HB distance, and *r*_*HB*_is the cutoff distance. While the details of our parametrization approach for *U*_stack_ and *U*_*HB*_ interactions are provided in the results and discussion section, the optimized parameters for *U*_stack_and *U*_*HB*_interactions can be found in **Table 2 and 3**, respectively. The choice of the cutoff distance for both the stacking and hydrogen bonding potentials ensures that the value of the 12-10 LJ potential becomes negligible beyond that distance.

**Table 2:** Non-bonded interaction parameters for stacking interactions associated with eq. 4.

| Base Pairs | Stacking |  | Base Pairs | Stacking |  |
| --- | --- | --- | --- | --- | --- |
| | $\sigma_{stack}$ (Å) | $\delta_{stack}$ (kcal/mol) | | $\sigma_{stack}$ (Å) | $\delta_{stack}$ (kcal/mol) |
| A-A | 3.6 | 9.7 | T-C | 3.6 | 6.9 |
| A-T | 3.6 | 8.6 | T-G | 3.6 | 9.2 |
| A-C | 3.6 | 7.8 | C-C | 3.6 | 6.2 |
| A-G | 3.6 | 10.3 | C-G | 3.6 | 8.3 |
| T-T | 3.6 | 7.7 | G-G | 3.6 | 11.0 |
| $r_{stack}$ (Å) | 6.2 | | | | |

**Table 3:** Non-bonded interaction parameters for hydrogen bonding interactions associated with eq. 5.

| Watson Crick Base Pairs | Hydrogen Bonding |  |
| --- | --- | --- |
| | $\sigma_{HB}$ (Å) | $\delta_{HB}$ (kcal/mol) |
| A-T | 6.0 | 2.7 |
| C-G | 5.5 | 3.3 |
| $r_{HB}$ (Å) | 9.5 | |

The *U*_*excluded*_ term accounts for excluded-volume interactions between two types of pairs: (a) base beads on the same strand that are one base position apart (i.e., *i, i*+2 pair), and (b) bases that do not contribute to the *U*_stack_and *U*_*HB*_potentials. These excluded-volume interactions are represented by the purely repulsive Weeks-Chandler-Andersen (WCA) potential (51):

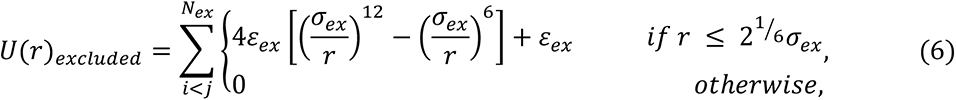

where *N*_*ex*_ represents the number of excluded-volume interaction pairs, *ε*_*ex*_ corresponds to the interaction strength, and *σ*_*ex*_ denotes the excluded-volume diameter. For all the beads, these parameters are set as: *ε*_*ex*_ = 4 kcal/mol and *σ*_*ex*_ = 5.5 Å.

The *U*_*electrostatics*_ term accounts for the electrostatic interactions between sugar-phosphate backbone beads, which carry an overall charge of -1. These electrostatic interactions are represented by the Debye-Hückel (DH) potential (52), which is applicable for low-salt concentrations commonly found in biological systems under physiological conditions. The DH potential is given by:

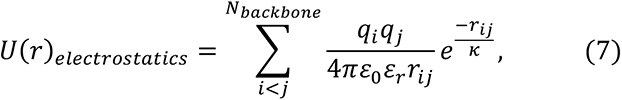

where *N*_*backbone*_ represents the number of backbone pairs, *q_i_* is the charge of the backbone bead, *ε*_3_ corresponds to the vacuum dielectric permittivity, *ε*_*r*_ denotes the relative dielectric constant of water (set to 80), and *κ* is the Debye screening length, which is taken as 10 Å and 8.8 Å for 100 mM and 120 mM salt concentrations, respectively. To accelerate the computational efficiency, a cutoff distance of 3.5*κ* is employed.

#### 1.1.1. Improving the directionality of the hydrogen bonding interactions

In order to address the isotropic nature of the non-bonded interactions in the 2-bead CG DNA model, which may not accurately capture the directionality of hydrogen bonds in A-T and C-G base pairs, we propose an extension of our model called the 3-bead CG DNA model. This modification, inspired by the approach of Jayaraman and co-workers (53,54) to improve the directionality of hydrogen bonding interactions, can result in improved structural properties of dsDNA duplex. The 3-bead CG model is developed by introducing a small dummy bead onto each base bead, as shown in **Figure 2**. These additional beads, designated as a, t, c, and g for the respective bases (A, T, C, G), serve as intra- and inter-strand hydrogen bonding sites.

**Figure 2.**
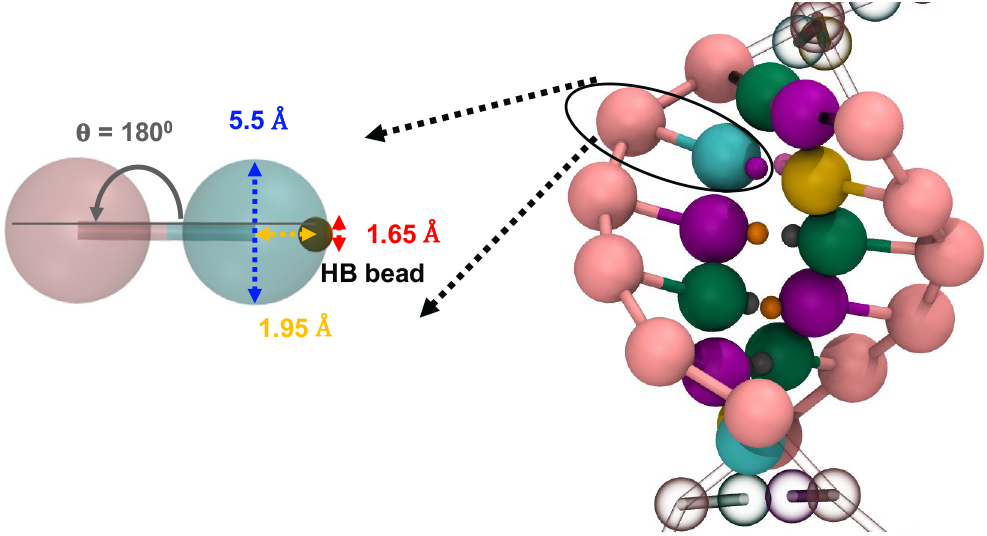
Schematic representation (not drawn to scale) of the 3-bead CG model of DNA, an extension of the 2-bead CG model (**Figure 1**), incorporating a small dummy bead on the base bead to capture the effective directionality of hydrogen bonding interactions. The model includes a sugar-phosphate backbone bead (pink color), bases A, T, C, G represented by green, violet, yellow, and cyan colors respectively, and HB beads a, t, c, g represented by grey, orange, purple, and brown colors respectively.

Previously, Jayaraman and co-workers (53,54) have demonstrated that by carefully adjusting the size and placement of such beads relative to their parent beads, effective directional interaction can be achieved. Following their approach, we set the size of the HB beads to 0.3 times the size of the base beads. The HB beads are positioned at an equilibrium bond distance of 1.95 Å from the centers of the base beads, using harmonic springs with a force constant of 50 kcal/mol. This arrangement allows the HB bead to be partially embedded within the base bead, while exposing it partially to enable effective directional interactions. The mass of the HB beads is assigned as 0.5 times the mass of the base bead. To further restrict the movement of these small HB beads in relation to their parent base beads, we introduce an angle potential involving the HB bead, its parent base bead, and the corresponding backbone bead. This angle potential is described by the cosine/squared-style harmonic potential (eq. 3), with *K*_*angle*_ set to 80 kcal/mol and *θ*_0_ set to 180 degrees.

Since the HB beads are introduced solely to facilitate hydrogen bonding interactions, pairwise non-bonded interactions involving these HB beads are defined only for a-t and c-g HB pairs, using the 12-10 LJ form (eq. 5). The interaction cutoff is set to 2 *σ*_*HB*_for these interactions. The parameters for the *U*_*HB*_ interactions are summarized in **Table 4**. Additionally, similar to the 2-bead model, we exclude hydrogen bonding interactions between adjacent HB beads (*i*, *i*+1 pair) and next nearest neighbor HB beads (*i*, *i*+2 pair) on the same DNA strand to prevent unphysical base pairing within the strand and the formation of local ring-like structures.

**Table 4:**
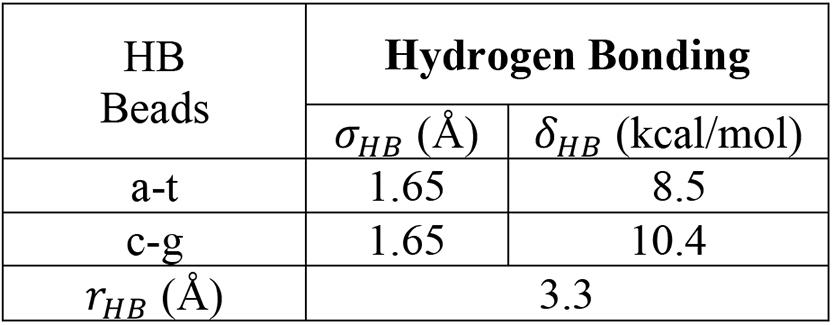
Parameters for hydrogen bonding interactions between small HB beads associated with eq. 5.

| HB Beads | Hydrogen Bonding |  |
| --- | --- | --- |
| | $\sigma_{HB}$ (Å) | $\delta_{HB}$ (kcal/mol) |
| a-t | 1.65 | 8.5 |
| c-g | 1.65 | 10.4 |
| $r_{HB}$ (Å) | 3.3 | |

### 1.2. Protein-DNA cross interactions

In previous studies, the determination of cross-interaction parameters for protein-DNA association has often focused on calibrating either short-range van der Waals (vdW) interactions or long-range electrostatic interactions to reproduce experimental measurements such as the molar dissociation constant (*K*_*d*_). For example, Takada and co-workers (29,43) considered electrostatic interactions as the dominant factor and calibrated the charge on the CG phosphate bead, while only including excluded volume interactions for short-range contacts, in order to reproduce experimental *K*_*d*_ values. On the other hand, Lebold and Best (28) parameterized the ɛ parameter of the Gō-type potential function to match experimental *K*_*d*_values. However, these approaches are partially dependent on the specific protein-DNA systems, which limits the transferability of cross-interaction parameters for studying sequence-dependent effects.

In this work, we incorporate both short-range vdW and long-range electrostatic interactions for protein-DNA interactions. However, instead of calibrating the parameters based on experimental *K*_*d*_ values, we utilize the all-atom hydropathy scale to define non-bonded protein-DNA interactions. Specifically, we draw the energy parameters for short-range vdW interactions between beads representing amino acids and nucleotides from the HPS modeling framework (48–50). The HPS CG protein model and DNA parameters in the HPS modeling framework are described in the following subsections.

#### 1.2.1. CG protein model

The previously developed CG model for proteins, known as HPS-Urry, has been successfully used to study the sequence-dependent LLPS of intrinsically disordered proteins (IDPs) (50). The HPS-Urry model represents each amino acid with a single bead, which is positioned at the Cα atom and connected to neighboring beads via harmonic spring. The bonds between adjacent beads are described by a harmonic potential (eq. 2) with equilibrium bond length, *r*_0_, and force constant, *K*_*bond*_, set to 3.8 Å and 10 kcal/mol/ Å^4^, respectively. For the HPS-Urry model, long-range electrostatic interactions are modeled using eq. 7, while short-range vdW interactions are represented by a modified LJ potential that allows for independent scaling of attraction and short-range repulsion between two residues, denoted as *i* and *j* (55),

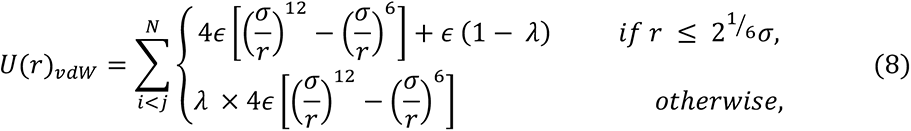

where *N* represents the number of residue pairs, *λ* corresponds to the average hydropathy, *σ* denotes the average diameter, and *ϵ* is the interaction strength between two residues, denoted as *i* and *j.* In this equation, the value of *ϵ* is set to 0.2 kcal/mol, and the hydropathy (*λ*) and vdW diameter (*σ*) for a pair of residues are calculated using arithmetic mixing rules. It should be noted that among twenty amino acids, Arg and Lys have a positive charge (+1), while Asp and Glu have a negative charge (-1). Further details about the HPS-Urry model can be found in the paper by Regy et al. (50).

#### 1.2.2. CG DNA-Protein interaction parameters in the HPS modeling framework

The hydropathy scale of DNA beads is determined based on the HPS modeling framework of an all-atom force field that assigns each atom as hydrophilic or hydrophobic depending on their partial charges, similar to the Kapcha-Rossky (KR) hydropathy scale of amino acids (56). In this work, we use the partial charges from the CHARMM27 all-atom force field to obtain the hydropathy values for each nucleotide bead. These parameters are summarized in **Table 5**. Similar to the protein-protein interactions, in eq. 8 *ϵ* is set to 0.2 kcal/mol for DNA-protein interactions. The hydropathy and vdW diameter values for a pair of amino acids and nucleotides are determined using arithmetic mixing rules. It is important to note that for the 3-bead model, the cross interactions between HB beads and protein residues have *ϵ* = 0, ensuring that the dummy HB beads do not interact with the CG protein beads.

### 1.3. Simulation details

#### 1.3.1. Parallel tempering simulations

In this subsection, we outline the system and simulations performed to parameterize and validate the 2-bead and 3-bead CG DNA models. We choose to parameterize the model at 120 mM salt concentration using a 14 bp oligomer dsDNA (S1: 5‘-GCGTCATACAGTGC-3’ and its complement S2: 5‘-GCACTGTATGACGC-3’) since both computational and experimental melting data are available for this DNA duplex (57–59).

To investigate the melting behavior of the CG DNA models, we employ parallel tempering simulations (60). Specifically, we perform replica exchange molecular dynamics simulations (REMD) with 32 replicas over a temperature range of 250–450K using the LAMMPS package (Oct 2020 version) (61). We implemented both 2-bead and 3-bead CG DNA models in LAMMPS for these simulations. The CG simulations are conducted using Langevin dynamics in an NVT ensemble, where the temperature is controlled by a Langevin thermostat with a damping parameter (referred to as ‘damp’ in LAMMPS package) set to 1000 time steps.

At the beginning of each simulation, the 14 bp dsDNA is randomly placed in a cubic simulation box with dimensions of 300 Å and periodic boundary conditions are applied in the x, y, and z directions. Each replica is simulated for 0.5 *μ*s, resulting in a total simulation time of 16 *μ*s. A time step of 10 fs is used, replica swaps are attempted every 100 steps, and configurations are sampled every 50,000 time steps. The first 50 ns of simulation data are discarded as equilibration, and the remaining 0.45 *μ*s trajectory is used for analysis. The melting curve and structural properties are computed by averaging over a total of 900 configurations from each replica.

#### 1.3.2. Umbrella sampling simulations

In this subsection, we outline the system and simulations performed to validate the protein-DNA cross interactions. Inspired by the work of Lebold and Best (28), we focus on computing the molar dissociation constant (*K*_*d*_) between a model protein-DNA system consisting of the C-terminus of histone H1 protein chain and a 20-bp dsDNA (see **SI Tables S1 and S2**). The disordered C-terminus of histone H1 comprises of 111 residues, with a net charge of +43 due to the presence of 45 positively charged and 2 negatively charged residues, while 20-bp dsDNA backbone carries a net charge of -40.

To determine the radially averaged potential of mean force (PMF) between the center of masses (COM) of the protein and DNA chains, we employ umbrella sampling with replica exchange. These simulations are conducted using the LAMMPS package (Oct 2020 version) (61), that incorporates the CG HPS-Urry protein model and the 2-bead/3-bead CG DNA models, augmented with SSAGES (62). To set up the system, we generate an initial configuration where the two chains are placed in a large cubic box with dimensions of 100 nm, ensuring that the COMs of the chains are 5 Å apart. This initial configuration is then subjected to steepest descent energy minimization to remove overlaps.

The umbrella sampling simulations are performed with a total of 40 replicas, with the umbrella potential set to 250 kJ mol^-1^ nm^-2^ (or 0.5975 kcal/mol/Å^2^) between the COMs of the protein and DNA chains. Each umbrella is simulated for 0.5 *μ*s, resulting in a total simulation time of 20 *μ*s, at a temperature of 300 K and 100 mM salt concentration. The CG simulations employ Langevin dynamics in an NVT ensemble, with the temperature maintained using a Langevin thermostat with a damping parameter of 1000 time steps. The weighted histogram analysis method (WHAM) (63,64) is utilized to obtain the free energies, which is then corrected with the missing Jacobian contribution to obtain the PMF.

The resulting PMF is further used to calculate *K*_*d*_ using the following eq.:

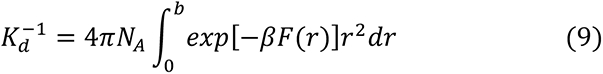

where *N*_A_ is Avogadro’s constant, b is the distance at which the PMF reaches its limiting value of 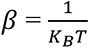 (where *K*_*B*_ is the Boltzmann constant and T the absolute temperature), *F*(*r*) is the PMF, and *r* is the intermolecular distance.

#### 1.3.3. Large-scale Langevin dynamics simulations of nucleosomes and LLPS

This subsection outlines the systems and simulations performed to demonstrate the suitability of our CG DNA models for facilitating simulations with molecular resolution in biological applications. First, we simulate a mono-nucleosome on a time scale of several μs to generate extensive residue-level conformational ensembles of DNA and histone tails. Next, we simulate this nucleosome with HP1α homodimers to gain mechanistic insights into the effects of protein-protein and protein-DNA interactions on the LLPS of HP1α in the presence of the nucleosome.

Nucleosomes are fundamental subunits of chromatin structure, consisting of a histone octamer composed of four types of core histones (H3, H4, H2A, and H2B), with two copies of each, and approximately 145-147 bp of DNA spooled around them (65). All core histones have disordered, positively charged N-terminal tails followed by small histone-fold domains. Additionally, H2A has a disordered C-terminal tail. To accurately represent the histone proteins with folded and disordered domains, as well as the nucleosomal DNA, we construct the initial CG structure of the nucleosome using the all-atom model from Peng et al. (33). Specifically, we use the nucleosome structure designated as Model A in their paper, which is constructed using PDB IDs 1AOI and 1KX5 as a template and includes a 20 bp linker DNA flanking the histone core on the entry and exit sites (see **SI Tables S1 and S2** for sequence information). For the histones, we construct a single-bead representation of each amino acid using the all-atom Cα positions. For the DNA we construct a 2-bead representation using COM of multiple atom positions (**Figure 3a**). The definitions for the histone regions and the atom representations used for the 2-bead CG DNA are provided in **SI Tables S3 and S4**, respectively. Additionally, we simulate nucleosomes without histone tails, by simply removing the residues that are considered part of each disordered tail. For our purposes, similar to Panchenko and co-workers (33,34), we assign integer values to the DNA superhelical locations (SHL) to create a DNA coordinate frame and classify the spooled versus linker DNA, as shown in **Figure 3b**.

**Figure 3.**
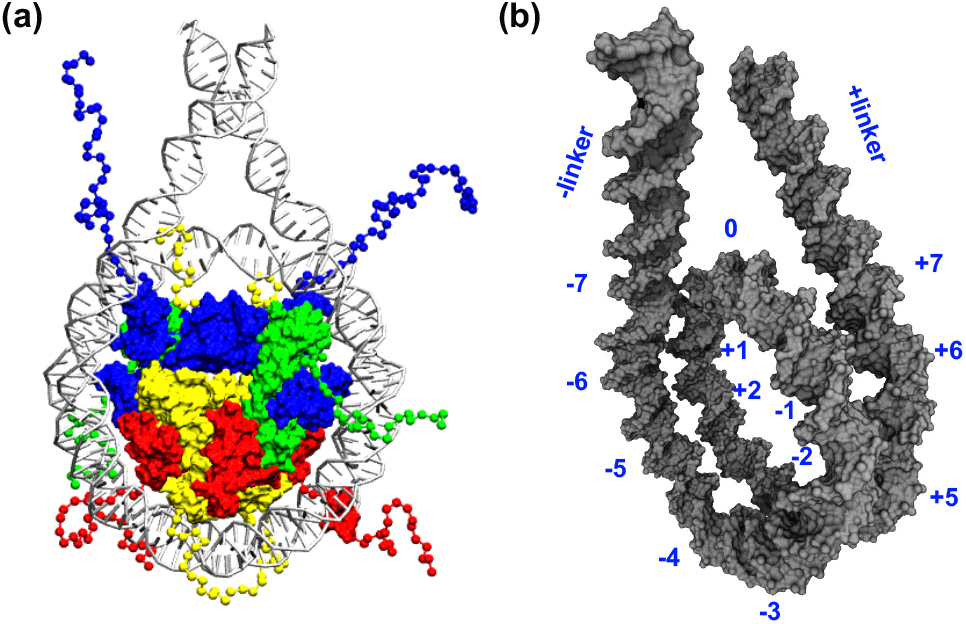
(a) Initial coarse-grained configuration of the nucleosome, and (b) The DNA coordinate frame: zero corresponds to the dyad position and integers represent the superhelical locations (SHL) of the nucleosomal DNA. We note that in the snapshot histones H3, H4, H2A and H2B are shown in blue, green, yellow, and red colors respectively, while DNA is shown in silver color, and for better representation, the histone-core shown as surface representation is created by overlaying atomistic folded helices of the histones over the coarse-grained rigid body beads.

Like histone proteins, HP1α is a multi-domain protein consisting of two highly conserved folded domains: the chromodomain (CD) and the chromoshadow domain (CSD), as well as three disordered regions: the N-terminal extension (NTE), the hinge region, and the C-terminal extension (CTE) (see **SI Table S1**). As we have done previously (66), we construct a single-bead representation of each amino acid using the all-atom Cα positions and represent HP1α homodimer by treating the CSD-CSD domains topologically together.

We perform CG MD simulations using the HOOMD-blue 2.9.7 package (67), augmented with azplugins (68), in which we have implemented both the HPS-Urry CG protein model and the 2- bead/3-bead CG DNA models. Similar to our previous work (66), we simulate the folded domains using rigid body dynamics by constraining the residues that are part of the rigid domain using the hoomd.md.constrain.rigid function (69).

To simulate the nucleosome, we first place the CG nucleosome structure we generated at the center of a large cubic box with dimensions of 100 nm. To relax the chains, we subject the initial CG configuration to steepest descent energy minimization. After this initialization stage, we run the simulation for 5 *μ*s at T = 300 K and 100 mM salt concentration. Similarly, for simulating the HP1α + Nucleosome system, we first generate an initial configuration by placing 50 chains of HP1α and nucleosome in a large cubic box with dimensions of 100 nm. To remove overlaps, we subject this initial configuration to steepest descent energy minimization. Next, to perform phase coexistence simulation, we resize this cubic box and create an initial slab configuration of dimensions 17x17x100 nm^3^. After this initialization stage, we run the simulation for 5 *μ*s at T = 320 K and 100 mM salt concentration, as done in our previous study (66).

We evolve the CG simulations using Langevin dynamics approach in an NVT ensemble, with a time step of 10 fs. The temperature is maintained using a Langevin thermostat with the friction factor *γ* set using 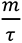, where *m* is mass of each CG bead and *r* is the damping factor set to 1000 ps.

Lastly, when showcasing the conformational ensembles and computing the concentration profiles and the intermolecular contact maps, we skip the initial 1 *μ*s trajectory as equilibration.

## 2. Results and Discussion

### 2.1. Parameterization strategy of the CG DNA models

Our objective is to replicate the melting thermodynamics and structural features of a 14 bp dsDNA sequence (S1S2) at 120 mM salt concentration. To obtain melting curves (fraction of melted DNA against temperature) from simulation data, we determine the hybridized or melted state of a DNA strand by counting the number of HB sites involved in hydrogen bonds with the complementary strand. A hydrogen bond is considered formed if the distance between HB sites is < 1.5*σ*_*HB*_. We classify a DNA strand as hybridized if at least half of its HB sites are engaged in hydrogen bonds. The temperature at which the fraction of the melted state reaches half on the melting curve is defined as the melting temperature (Tm). To determine the model parameters, we utilize an optimization scheme to find the parameters for both the stacking interactions and hydrogen bonding interactions which reproduce the melting curves of dsDNA, along with its structural features, while keeping all the other bonded and non-bonded parameters unchanged.

We carry out the parameterization for the 2-bead model in the following manner. We choose the initial values for stacking and hydrogen bonding interactions based upon one of our previous works, where we obtained PMFs for DNA base pairs as a function of distance for base pair stacking and base pair hydrogen bonding free energies (70). We assign initial values for hydrogen bonding interactions between A-T pairs and C-G pairs based on Table 1 of the reference (70). Similarly, the initial values for stacking interactions between each unique base pair combination (A-A, T-T, C-C, G-G) are obtained from Table 1 of the same reference (70). We use Lorentz-Berthelot mixing rules to determine the parameters for stacking interactions involving cross base pairs (A-T, A-C, A-G, T-C, T-G, C-G).

We systematically scale only the energy parameters for both the stacking interactions and hydrogen bonding interactions to reproduce the experimental melting transition, along with the structural features of dsDNA. It is important to note that during parameterization, we keep the size parameters for both stacking and hydrogen bonding interactions, as well as the relative strength of stacking interaction among all the 10 pair combinations (i.e., A-A, A-T, … G-G) and the relative strength of hydrogen bonding interactions between A-T and C-G pair unchanged. We define the scaling parameters for stacking and hydrogen bonding energy parameters as Δ_stack_and Δ_*HB*_, respectively. In **Supplementary Information (SI) Figure S1**, we demonstrate how changes in Δ_stack_ and Δ_*HB*_ influence the melting curves for select cases. The final parameters, summarized in **Table 2 and 3,** have values of 2.1 for Δ_stack_ and 0.95 for Δ_*HB*_, respectively.

Similarly, for the 3-bead model, we re-parameterize the hydrogen bonding energies to match the melting thermodynamics of the dsDNA duplex. The final parameters, summarized in **Table 5**, have values 3.1 times that of the value of *δ*_*HB*_ in **Table 3**.

**Table 5:** Short-range vdW energy parameters for nucleotides in the HPS framework associated with eq. 8.

| CG Bead | $\lambda$ value |
| --- | --- |
| backbone | 0.38 |
| ADE | 0.40 |
| THY | 0.54 |
| CYT | 0.59 |
| GUA | 0.35 |

### 3.2 Melting behavior of dsDNA

In this section, we examine the melting behavior of the 14 bp dsDNA sequence (S1S2) at 120 mM salt concentration. **Figure 4** presents the melting curves for both 2-bead and 3-bead CG models, utilizing the optimized parameters outlined in **Tables 2-4**. To provide an additional measure of Tm independent of the specific pair cutoff distance used in the structure-based definition, we also calculate the heat capacity as a function of temperature (**Figure 4)**, and identify Tm from the heat capacity maximum, in accordance with the calorimetric definition (71).

**Figure 4.**
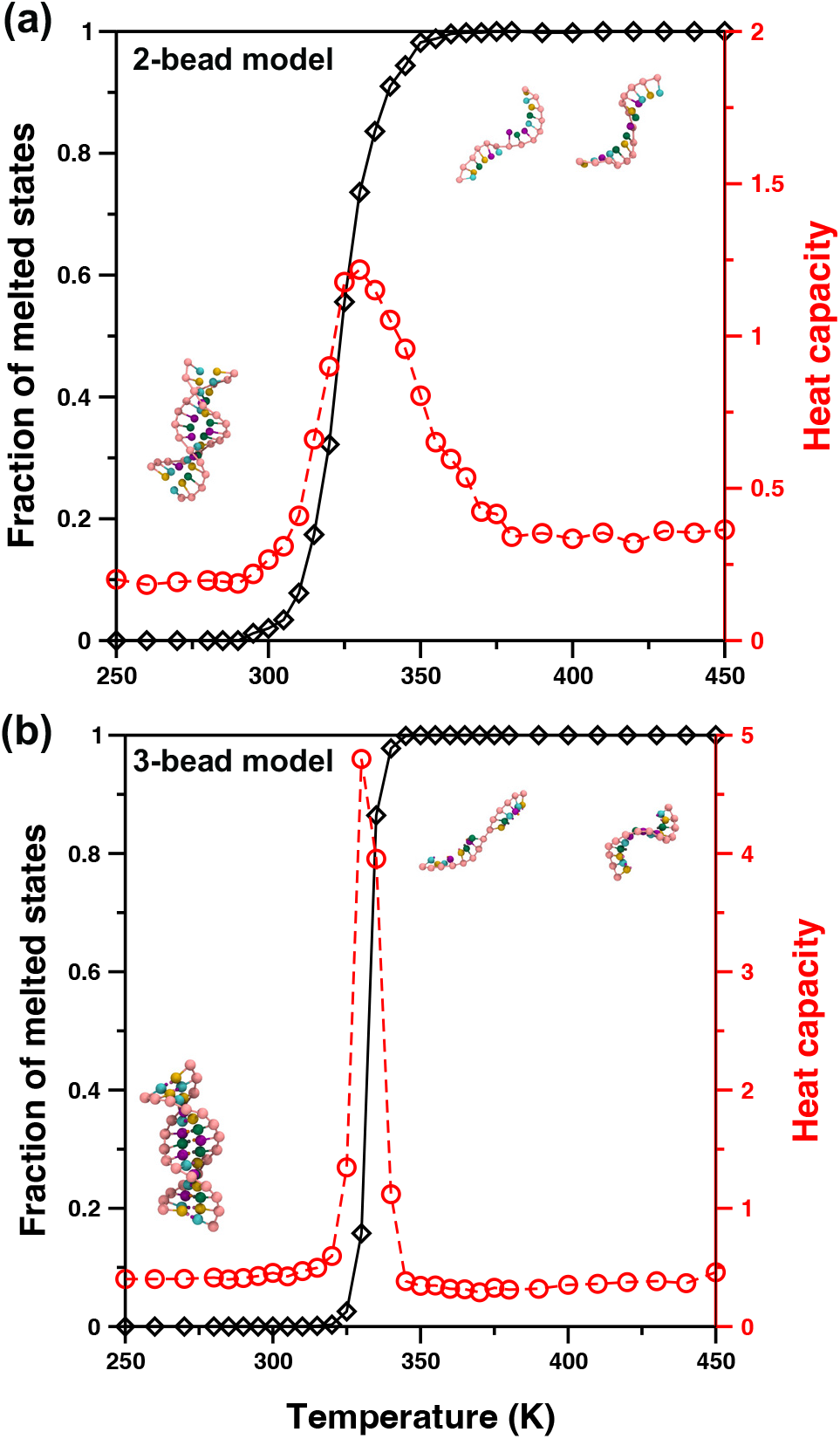
Thermodynamic melting behavior obtained from parallel-tempering simulations. The fraction of melted states (black symbols) and heat capacity (red symbols) as a function of temperature, and typical dsDNA configurations are shown for a pair of dsDNA S1S2: 5‘- GCGTCATACAGTGC-3’ obtained using our (a) 2-bead CG DNA model and (b) 3-bead CG DNA model. In the snapshots the sugar-phosphate backbone bead is shown in pink color, bases A, T, C, G are shown in green, violet, yellow and cyan colors respectively, while the HB beads a, t, c, g are shown in grey, orange, purple and brown colors respectively.

As anticipated, our observations reveal that at extremely low temperatures, nearly all states exhibit hybridization, while at high temperatures, all states become melted. The melting temperature from the 2-bead CG model is measured to be 329.3 ± 2.8 K, whereas the 3-bead CG model yields a melting temperature of 332.4 ± 1.2 K. As the parameters of both models are optimized to reproduce the experimental melting behavior of the same dsDNA at 120 mM, the melting temperatures obtained from simulations are, by design, in excellent agreement with the experimentally determined melting temperature of 333.2 ± 0.5 K (59). Furthermore, our results demonstrate a close agreement in the temperature range over which melting occurs, which resembles the expected thermodynamic melting behavior observed in experiments (57,59).

### 3.3. Structural properties of dsDNA

To assess the ability of our CG DNA models to capture the local structural properties of hybridized dsDNA strands, we perform REMD simulations of a 32 bp dsDNA sequence: 5‘-ATACAAAGGTGCGAGGTTTCTATGCTCCCACG-3’ at T = 290 K (well below the melting temperature of ∼ 345 K) and 100 mM salt concentration. We focus on calculating several structural properties, including the helical width of the duplex, the number of base pairs per turn, the base rise, and the widths of the major and minor grooves. To approximate the helical axis, we employ the method described in reference (72). We select tetrads of nucleotides separated by 3 nucleotides, ensuring that the nucleotides are located at least 3 bases away from the termini to minimize end effects, as done by Hinckley et al. (22). Using this approximated helical axis, we determine the width of the duplex, the number of base pairs per turn, and the base rise, as outlined in the reference (72).

Furthermore, we evaluate the widths of the major and minor grooves associated with a specific base pair step using the method presented in reference (73). Specifically, we compute the widths of the major and minor grooves using the “TC” base pair step found at the 19^th^ step of the 32 bp sequence (5‘-ATACAAAGGTGCGAGGTT**<u>TC</u>**TATGCTCCCACG-3’) in order to mitigate any potential end effects.

The mean values and standard deviations of the local structural properties obtained from the simulations are presented in **Table 6**, along with the corresponding experimental data for comparison. The results indicate that the structure of dsDNA remains stable throughout the simulation, as expected. Notably, the inclusion of additional HB beads in the 3-bead model, which improved the effective directionality of the hydrogen bonding interactions among the bases, leads to enhanced agreement with the experimental data in terms of the local structural properties.

**Table 6:** Comparison of structural properties predicted by the 2-bead and 3-bead models to the values from the B-DNA crystal structure. Structural properties are obtained from the 32 base pair sequence 5‘-ATACAAAGGTGCGAGGTTTCTATGCTCCCACG-3’ at T = 290 K and 100 mM salt concentration. Experimental data at T = 293.15 K and 100 mM salt concentration are taken from references (22,74).

| Structural Property | 2-bead model | 3-bead model | Expt. |
| --- | --- | --- | --- |
| Base Rise (Å) | $3.0 \pm 0.3$ | $3.3 \pm 0.3$ | 3.4 |
| Helical Width (Å) | $23.8 \pm 0.7$ | $23.1 \pm 0.1$ | 23 |
| Base Pairs per Turn | $9.2 \pm 0.5$ | $9.9 \pm 0.2$ | 10 |
| Minor Groove Width (Å) | $15.6 \pm 0.7$ | $16.4 \pm 0.5$ | 17.1 |
| Major Groove Width (Å) | $13.9 \pm 0.4$ | $12.2 \pm 0.3$ | 11.8 |

While the values predicted using 2-bead CG DNA model exhibit slightly higher percentage error deviation from the experimental data compared to the 3-bead CG DNA model, overall, both models show good agreement with the experimental values for base rise, helix width, base per turn, and major/minor grooves.

### 3.4. Protein-DNA interactions

To demonstrate the general validity of the CG DNA models for studying protein-DNA systems, we compute the molar dissociation constant between the C-terminus of histone H1 protein and a 20-bp dsDNA (see **SI Tables S1 and S2**), where we utilize the HPS-Urry model for the protein along with both the 2-bead or 3-bead CG models for DNA. **Figure 5** presents the PMF profiles between the protein-DNA COMs, obtained from umbrella sampling simulations at T = 300 K and 100 mM salt concentration. Additionally, we provide plots of the COM versus time and histograms per replica in **SI Figure S2** to illustrate the sampled regions in each replica and the sufficient overlap between adjacent replicas, respectively.

**Figure 5.**
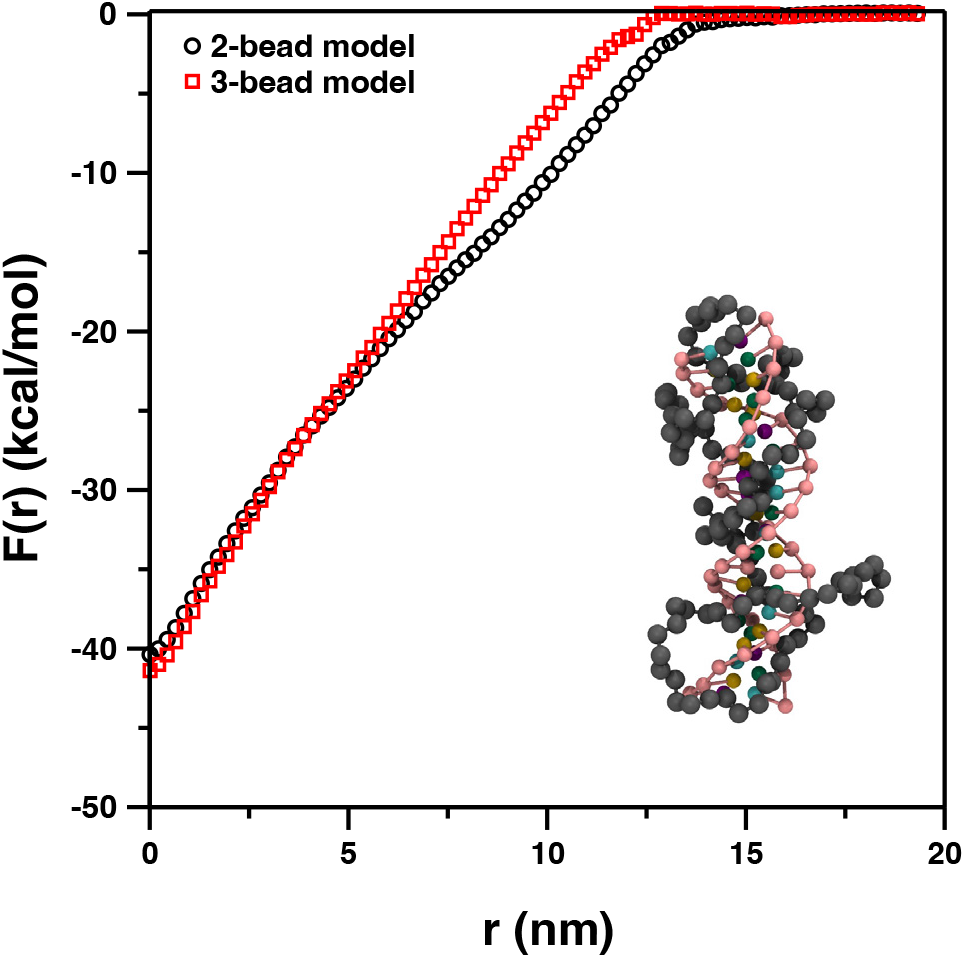
Representative potential of mean force, F(r), as a function of distance, between the center of mass of C-terminus of histone H1 protein and the center of mass of DNA molecules. The data is obtained from umbrella sampling simulations using both the 2-bead and 3-bead CG DNA models. Also, included are coarse-grained representations of the C-terminus of histone H1 protein and DNA (2-bead model). In the snapshot the sugar-phosphate backbone bead is shown in pink color, while the bases A, T, C, G are shown in green, violet, yellow and cyan colors, respectively. The protein beads are represented in gray color.

We find that the PMFs obtained using the 2-bead and 3-bead CG DNA models exhibit very similar profiles, as one would expect. The minor differences in PMFs are likely due to variations in the structure of the DNA duplex obtained from the 2-bead and 3-bead models. Using these PMFs in eq. 9, we also calculate the corresponding *K*_*d*_ values as 73.4 nM and 75.6 nM for the 2-bead and 3-bead CG DNA models, respectively, which are comparable to the experimental *K*_*d*_ for this system reported as 101 ± 20 nM under ambient conditions and 160 mM salt concentration (75), indicating that both 2-bead and 3-bead CG DNA models capture the protein-DNA intermolecular interactions reasonably well.

### 3.5. Large-scale protein-DNA simulations

The findings above indicate that both the 2-bead and 3-bead CG DNA models effectively capture the structural properties, thermodynamics of dsDNA hybridization, and protein-DNA interactions. Building on these results, we now aim to demonstrate the suitability of our model for large-scale simulations at the micron-scale, with molecular resolution, in various biological applications. In this regard, we utilize the 2-bead CG DNA model in combination with the HPS-Urry CG protein model and simulate a complete nucleosome. We investigate the nucleosomes, aiming to gain insights into the role of histone tails in modulating the conformational ensembles of DNA and LLPS of HP1α proteins.

#### 3.5.1. Role of histone tails in stability and unwrapping of nucleosomal DNA

In this section, we investigate the impact of histone tails on the behavior of nucleosomal DNA. We begin by comparing the MD trajectories of the nucleosome systems with and without histone tails to assess the influence of histone tails on the structure of nucleosomal DNA. Previously, both computational (33,34,43,45,76,77) and experimental (78–80) studies have demonstrated that histone tails play a significant role in restricting DNA breathing motions and unwrapping. Consistent with these findings, our observations reveal that nucleosomes with histone tails exhibit minimal DNA unwrapping, with transient detachment of a few base pairs at both entry and exit DNA sites. In addition, we observe that the tails of different histone types exhibit a preference for interacting with specific regions of DNA, suggesting that histone tail-DNA interactions play a role in restricting DNA unwrapping and breathing (**Figure 6a**). Conversely, when the histone tails are truncated, we observe spontaneous unwrapping of DNA from the core of histone octamer, with up to approximately 30 base pairs becoming unwrapped from the histone core at SHL ± 4 locations (**Figure 6b**, see **Figure 3b** for SHL DNA coordinate system).

**Figure 6.**
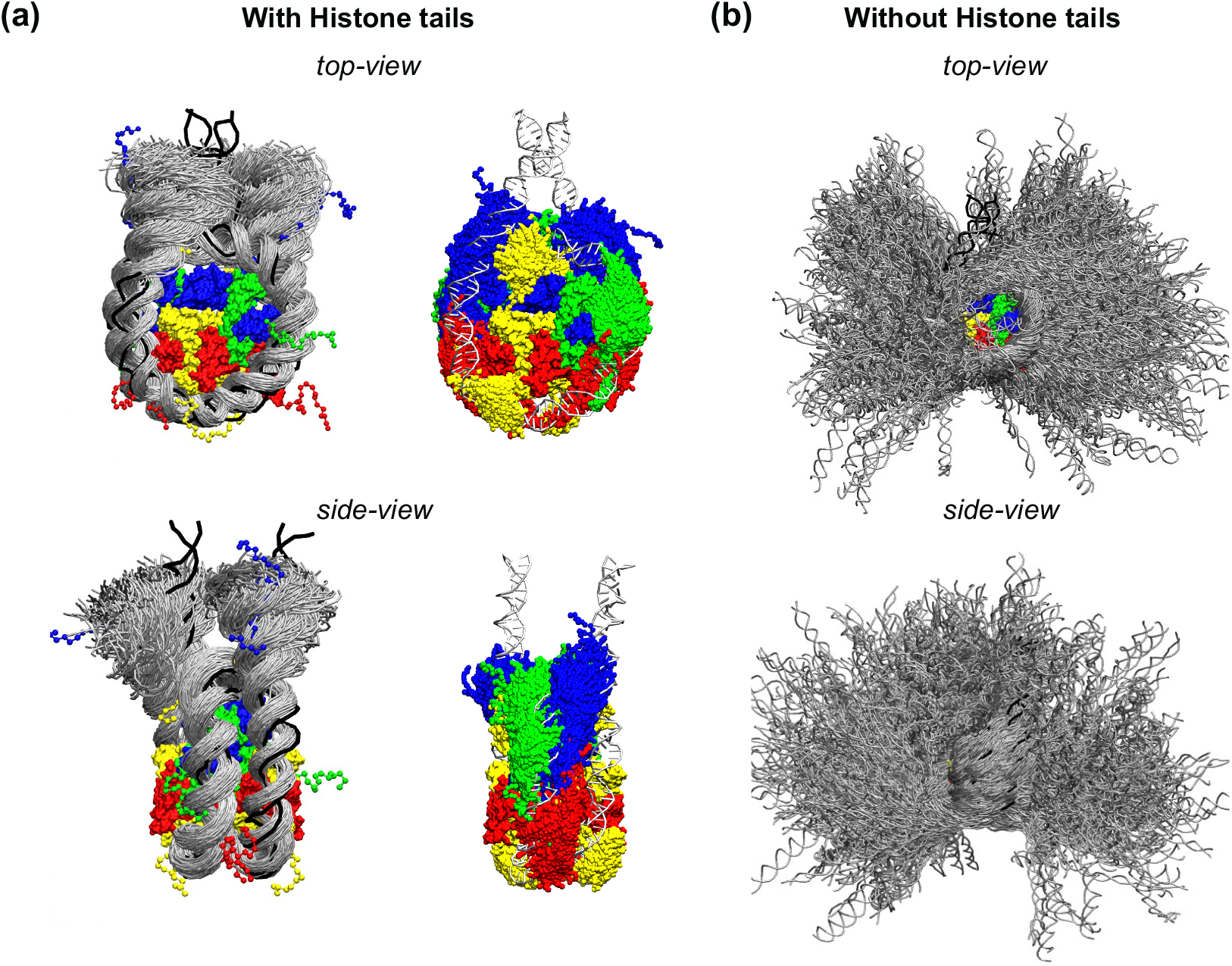
The conformational ensembles of (a) DNA (left) and histone tails (right), separately, and (b) DNA alone, for nucleosome systems with and without histone tails, respectively. These snapshots are generated by overlaying 1000 snapshots over the last 4 µs of the simulation trajectory. In the snapshots, the histones H3, H4, H2A and H2B are depicted in blue, green, yellow, and red colors, respectively, while the DNA is represented in silver color. The black lines indicate the initial configuration of the DNA at the beginning of the simulation. It is important to note that, for better visualization, the representation of the histone core as a surface is created by superimposing atomistic folded helices of the histones onto the CG rigid body beads. These snapshots are rendered using VMD software (81).

To further characterize this behavior, we first computed the intermolecular contact map between the histone octamer core and DNA, by analyzing vdW contacts averaged over the entire trajectory, and found that for nucleosome with histone tails, the probability of intermolecular contacts between the histone core and DNA remains relatively consistent across different SHLs. In contrast, for nucleosome without histone tails, the probability of intermolecular contacts between the histone core and DNA are significantly lower for SHL less than ± 4 locations (see SI **Figure S3**).

Next, we analyzed the molecular interactions between histone tails and DNA. **Figure 7** illustrates that different histone tail types preferentially interact with specific regions of the DNA (see also **SI Figure S4**). The H3 tails, being the longest among the histone tails, interact with multiple regions of the DNA, including near the dyad (SHL 0) and at SHL ±1, ±2, ±7, as well as with the linker DNA at the entry and exit sites. Similarly, the H4 tails, despite being the shortest, form a DNA-binding interface near the dyad and at SHL ±2. The H2B tails also interact with the DNA in multiple regions, specifically from SHL ±3 to ±6. The findings align with previous studies that characterized histone tail-DNA interactions (33,34,43,78,79).

**Figure 7.**
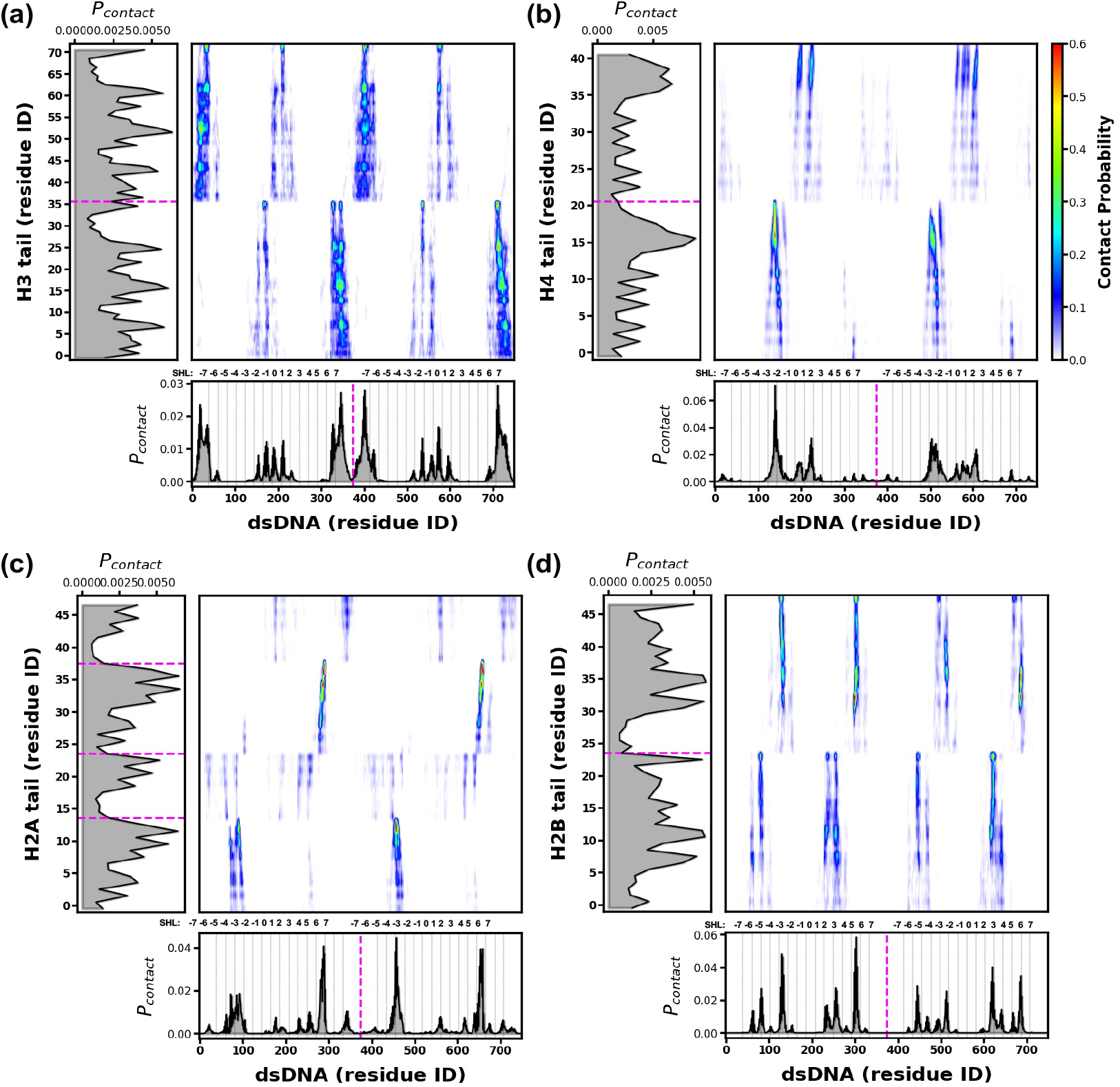
Intermolecular contacts between DNA and the tails of histones: (a) H3, (b) H4, (c) H2A, and (d) H2B. The preferential interactions are highlighted in red color. The dotted magenta color lines on the x-axis and y-axis represent the boundaries between the two DNA strands and the histone tails, respectively. Additionally, the gray color grid lines on the x-axis indicate the superhelical locations (SHL) in accordance with the SHL coordinate system used in this study (refer to **Figure 3b** for further details).

Contrary to the H3, H4 and H2B tails, the two tails of each H2A histone exhibit distinct interactions with DNA. One of the H2A tails predominantly binds to the nucleosomal DNA at SHL ±5, ±6, while its C-terminus spans all the SHL regions, including the linker DNA at the entry site. In contrast, the other H2A tail’s N-terminus primarily binds at SHL +5, while its C-terminus is mostly bound near the dyad and the linker DNA at the exit site. These observations suggest that the H2A histone tails are highly dynamic and encompass a more extensive DNA-binding interface, covering a larger area on the DNA compared to other tails. However, it is worth noting that these observations of H2A tail-DNA interactions slightly differ from previous reports. Peng et al. showed in their simulations that either of the H2A N-terminal tails bind to the nucleosomal DNA at SHL ± 4, whereas either of the H2A C-terminal tails primarily bind at SHL ± 7 and near the dyad (33). We attribute these differences to the limited timescale of the all-atom simulation or need for further improvements in the CG model.

In addition, consistent with previous observations (33,34), we find that the positively charged lysine/arginine-rich patches present on the histone tails mediate prominent interactions with DNA. These observations highlight the significance of interactions between the negatively charged sugar-phosphate backbone of DNA and RK/KR/KK residues of histone tails in controlling the DNA breathing process. While our findings align with previous simulation studies of nucleosomes, which characterized DNA unwrapping and breathing dynamics, it is important to note that our CG simulations can allow for more efficient sampling of conformational space, as our single 5 *μ*s CG simulation captures multiple instances of unwrapping and rewrapping events of the DNA at both entry and exit sites (see **SI Movie M1**).

Overall, our analysis of the dynamic MD ensemble of the nucleosome structure, in terms of DNA and histone tail conformations and histone tail-DNA intermolecular interactions, confirms that histone tails interact with specific regions of nucleosomal or linker DNA and have a direct impact on the DNA and nucleosome geometry. This provides insights into how histone-DNA interactions may regulate the accessibility of histone tails or DNA.

#### 3.5.2. Molecular insights into the role of histone tails in modulating LLPS of HP1α proteins

In order to further demonstrate the usefulness of the CG DNA models developed in this study, we conducted simulations to investigate the LLPS of HP1α proteins in the presence of a nucleosome. These simulations provide valuable insights into how the presence of nucleosomes affects the LLPS of HP1α and sheds light on the potential role of HP1α-nucleosome interactions in chromatin organization and compaction.

**Figure 8a** shows simulation snapshots and compare concentration profiles of HP1α in three different systems: pure HP1α, HP1α + free dsDNA, and HP1α + nucleosome at 320K. It is worth noting that in the simulation of HP1α + free dsDNA, we simulated a single chain of nucleosomal dsDNA along with 50 chains of HP1α. The figure illustrates that both the free dsDNA and the nucleosome gets incorporated into the HP1α droplet (see SI **Figure S5** for concentration profiles of nucleosome within the droplet). The concentration profiles of HP1α indicate that while the presence of free dsDNA promotes the LLPS of HP1α (lower dilute phase concentration), which has been known (5,66), intriguingly the influence of nucleosome on the LLPS of HP1α is less pronounced (similar dilute phase concentrations).

**Figure 8.**
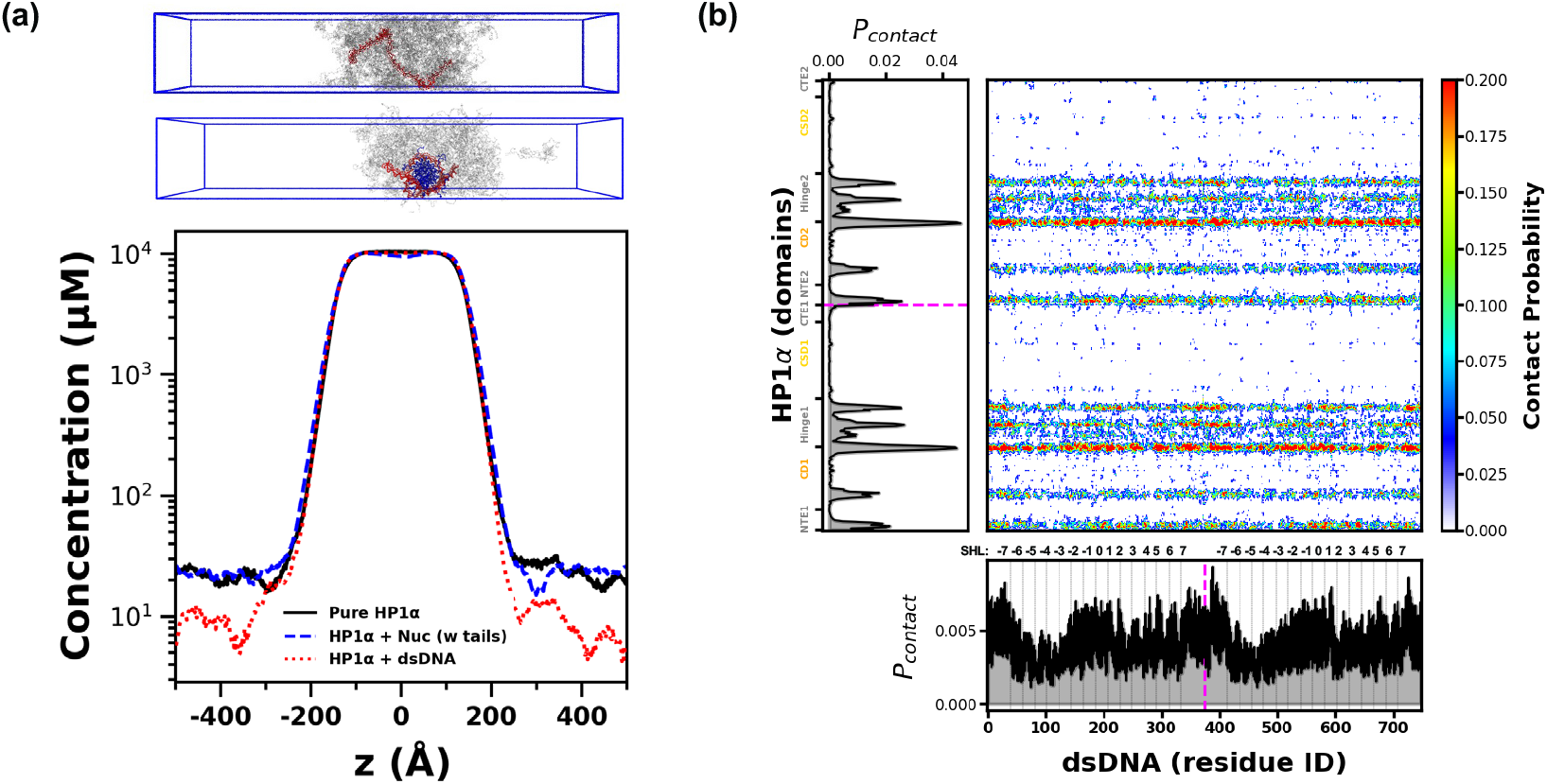
LLPS of HP1α with nucleosome. (a) Snapshots of phase coexistence slab simulations, and comparison of the concentration profiles of HP1α for different systems (see legend). The snapshots show HP1α condensates in gray color, while DNA and histones are depicted in red and blue colors, respectively. (b) Intermolecular contacts between nucleosomal DNA and HP1α. Preferential interactions are shown in red color. We also note that the dotted magenta color lines on the x-axis and y-axis indicate the demarcation between HP1α dimers and the two DNA strands, whereas the gray color grid lines on the x-axis show superhelical locations (SHL).

To understand this behavior and examine the interactions between HP1α and the nucleosome, we quantified the HP1α-DNA interactions by computing the intermolecular contact maps. **Figure 8b** shows that HP1α interacts with DNA through patches of positively charged lysine/arginine-rich regions in the hinge, NTE and CD region, which is consistent with previous experimental and computational studies (5,27,66,82). Interestingly, we observed that while HP1α form contacts with the entire DNA strand, the contact probabilities are generally higher for SHL near the dyad and at the DNA entry and exit sites, contrary to the SHL locations for histone core-DNA contacts (see **SI Figure S6a**). This suggests that histone tails, which interact favorably with DNA, compete with, and potentially antagonize the contacts between HP1α and DNA, thereby affecting the ability of DNA to promote LLPS of HP1α. To confirm that this behavior is not an artifact of kinetic limitation, we also performed simulations of HP1α with the nucleosome with a different initial configuration in which the nucleosomal dsDNA was not initially wrapped around the histone octamer. We observed that over the course of the simulation, the dsDNA started to interact with the histone proteins and eventually wrapped around the histone core (see **SI Movie M2**).

To understand how interaction of histone tails themselves gets affected when the nucleosome is partitioned inside the HP1α condensate, we computed the intermolecular contacts between histone tails and HP1α, as well as between histone tails and DNA. **SI Figures S7 and S8** show the respective intermolecular contacts. Our findings indicate that almost the entire length of the histone tails interacts with HP1α with similar propensity, without a specific arginine-/lysine-rich patch mediating the interactions between the histone tails and HP1α. Similar observations are made for histone core-HP1α contacts as well (see **SI Figure S6b**). This observation suggests that the binding of HP1α to the nucleosome is governed by diverse multivalent interactions, consistent with previous reports (6). Furthermore, we find that although histone tail-DNA interactions mediated by RK/KR/KK patches remain prominent, the tails lose their preferential ability to interact with specific DNA regions. In other words, interactions occur across all superhelical locations. This phenomena can likely be attributed to DNA sliding and nucleosome repositioning (42,43) as depicted in **SI Movies M1 and M2**). Consequently, our results suggest a significant increase in the dynamic behavior of DNA when the nucleosome partitions into the condensate of HP1α.

Overall, our findings demonstrate that our CG protein and DNA models effectively capture protein-DNA interactions, identify regions of high contact propensity, and offer valuable mechanistic insights. Specifically, we have shown that histone tails play a crucial role interacting with DNA, influencing its conformational ensemble, and modulating the interactions between HP1α and DNA. This, in turn, affects the ability of DNA to promote the LLPS of HP1α. Based on these observations, we propose that the regulation of nucleosome interactions with chromatin binding proteins and, consequently, epigenetic processes is largely governed by the modulation of DNA accessibility through histone tails. In our ongoing research, we are further utilizing these CG models to gain a comprehensive understanding of how nucleosome arrays recruit or impede interactions with specific regulatory proteins. Additionally, we aim to investigate how the LLPS of such regulatory proteins influences chromatin organization. These endeavors will provide valuable mechanistic insights into the dynamic interplay between nucleosomes, DNA, and regulatory proteins in the context of chromatin function and regulation.

## 3. Conclusions and Outlook

In this study, we have introduced predictive CG models for DNA, that balances molecular-level detail with reduced complexity by representing each nucleotide with two (or three) interaction sites. Our model incorporated isotropic potentials for base pair stacking and hydrogen bonding interactions, and remarkably, it accurately reproduces experimental measurements such as dsDNA melting temperatures and local structural properties of dsDNA, including duplex width, base rise, and major/minor groove widths. Moreover, our CG DNA model is compatible with the HPS CG protein models, as the non-bonded protein-DNA interactions are defined using the all-atom hydropathy scale.

To demonstrate the capabilities of our model in enabling large-scale protein-DNA simulations with molecular resolution, we conducted simulations of a nucleosome, with and without histone tails, and investigated the influence of histone tails on the conformational ensembles of DNA and the liquid-liquid phase separation of HP1α. By examining the interaction landscape of histone tails with DNA and HP1α proteins, we confirmed the important role played by histone tails in modulating DNA behavior and preventing spontaneous DNA unwrapping, which was observed in nucleosome systems with truncated histone tails. These findings further revealed the impact of histone tails on the propensity for liquid-liquid phase separation propensity of HP1α proteins.

Given that our CG DNA model incorporates both sequence information and grooving, it holds significant potential in various areas of computational biophysics. It can be applied to investigate the mechanisms of DNA hybridization, protein-DNA binding, nucleosome modeling, and explore the origins of binding affinities between proteins and specific DNA sequences. Additionally, the model can facilitate liquid-liquid phase separation of proteins with DNA and other related studies.

## Supplementary Materials

Additional CG model and simulation details, sequences of CH1 histone, and H3, H4, H2A, and H2B histone tails, and results for the extent of intermolecular interaction between DNA, histone core, histone tails, and HP1α, when the nucleosome is with and without histone tails and when the nucleosome is partitioned into the HP1α droplet.

## Acknowledgments

This material is based upon the work supported by the National Institutes of Health grant R01GM136917. Y.C.K. is supported by the Office of Naval Research via the U.S. Naval Research Laboratory base program. The computing resources were provided by Texas A&M High Performance Research Computing, specifically GRACE supercomputing cluster.

## Data and Software Availability

The source code required to run the 2-bead and 3-bead CG DNA models within the LAMMPS (Oct 2020) package is provided at the following location (https://github.com/utkarsk/CG-DNA-model). The repository also contains example input files to run simulations for a DNA duplex.

Other source data can be obtained from the corresponding author upon reasonable request.

