## Supplementary Information for "A coarse-grained DNA model to study protein-DNA interactions and liquid-liquid phase separation"

***Table S1.*** *Protein sequences used in this work*

| Protein | Sequence |
|---|---|
| **Histone H1 C-terminus tail** | KPGEVKEKAPKKKASAAKPKKPAAKKPAAAAKKPKKAV AVKKSPKKAKKPAASATKKSAKSPKKVTKAVKPKKAVA AKSPAKAKAVKPKAAKPKAAKPKAAKAKKAAAKKK |
| **Histone H3** | ARTKQTARKSTGGKAPRKQLATKAARKSAPATGGVKKP HRYRPGTVALREIRRYQKSTELLIRKLPFQRLVREIAQDFK TDLRFQSSAVMALQEASEAYLVALFEDTNLCAIHAKRVTI MPKDIQLARRIRGERA |
| **H3 tail, residues 1–36** | ARTKQTARKSTGGKAPRKQLATKAARKSAPATGGVK |
| **Histone H4** | SGRGKGGKGLGKGGAKRHRKVLRDNIQGITKPAIRRLAR RGGVKRISGLIYEETRGVLKVFLENVIRDAVTYTEHAKRK TVTAMDVVYALKRQGRTLYGFGG |
| **H4 tail, residues 1–21** | SGRGKGGKGLGKGGAKRHRKV |
| **Histone H2A** | SGRGKQGGKTRAKAKTRSSRAGLQFPVGRVHRLLRKGN YAERVGAGAPVYLAAVLEYLTAEILELAGNAARDNKKT RIIPRHLQLAVRNDEELNKLLGRVTIAQGGVLPNIQSVLLP KKTESSKSKSK |
| **H2A N-terminus tail, residues 1–13** | SGRGKQGGKTRAKA |
| **H2A C-terminus tail, residues 119–128** | KTESSKSKSK |
| **Histone H2B** | AKSAPAPKKGSKKAVTKTQKKDGKKRRKTRKESYAIYV YKVLKQVHPDTGISSKAMSIMNSFVNDVFERIAGEASRLA HYNKRSTITSREIQTAVRLLLPGELAKHAVSEGTKAVTKY TSAK |
| **H2B tail, residues 1–23** | AKSAPAPKKGSKKAVTKTQKKDGK |
| **HP1α (wild type)** | MGKKTKRTADSSSSEDEEEYVVEKVLDRRVVKGQVEYL LKWKGFSEEHNTWEPEKNLDCPELISEFMKKYKKMKEG ENNKPREKSESNKRKSNFSNSADDIKSKKKREQSNDIARG FERGLEPEKIIGATDSCGDLMFLMKWKDTDEADLVLAKE ANVKCPQIVIAFYEERLTWHAYPEDAENKEKETAKS |

***Table S2.*** *DNA sequences used in this work*

| DNA | Sequence |
|---|---|
| **14-bp DNA** | 5′-GCGTCATACAGTGC-3′ |
| **20-bp DNA** | 5′-ACGCCTGAAGAGTCTGGTGA-3′ |
| **32-bp DNA** | 5′-ATACAAAGGTGCGAGGTTTCTATGCTCCCACG-3′ |
| **Nucleosomal DNA** | 5′-<br>GGCGGCGGCTCGGCCAGTACTCCCGGCCCCCGCCATTTCGGA<br>CTGGGAGCGAGCGCGGCGCAGGCACTGAAGGCGGCGGCGGG<br>GCCAGAGGCTCAGCGGCTCCCAGGTGCGGGAGAGAGGTACG<br>GAGCGGACCACCCCTCCTGGGCCCCTGCCCGGGTCCCGACCC<br>TCTTTGCCGGCGCCGGGCGGG-3′ |

***Table S3.*** Definitions of histone alpha-helical core and tail regions

| Histone regions | Definitions |
|---|---|
| **Alpha helices in histone core** | **H3:** residues 45 to 56, 64 to 78, 86 to 114, 121 to 131<br>**H4:** residues 31 to 41, 49 to 76, 83 to 93<br>**H2A:** residues 17 to 21, 27 to 37, 45 to 73, 80 to 89, 91 to 97<br>**H2B:** residues 34 to 45, 53 to 81, 88 to 98, 101 to 118 |
| **H3 tail** | Residues 1 to 36 |
| **H4 tail** | Residues 1 to 21 |
| **H2A N- and C-terminus tail** | Residues 1 to 13 and 119 to 128 |
| **H2B tail** | Residues 1 to 23 |

***Table S4.*** Atom definitions used to generate initial position of the 2-bead CG nucleosomal DNA

| CG Beads | Atom names |
|---|---|
| **Backbone** | C1', C2', C3', C4', O4' |
| **ADE** | N1, C2, N3, C4, C5, C6, N6, N7, C8, N9 |
| **THY** | N1, C2, N3, C4, C5, C6, C7, O2, O4 |
| **CYT** | N1, C2, N3, C4, C5, C6, O2, N4 |
| **GUA** | N1, C2, N3, C4, C5, C6, N2, O6, N7, C8, N9 |

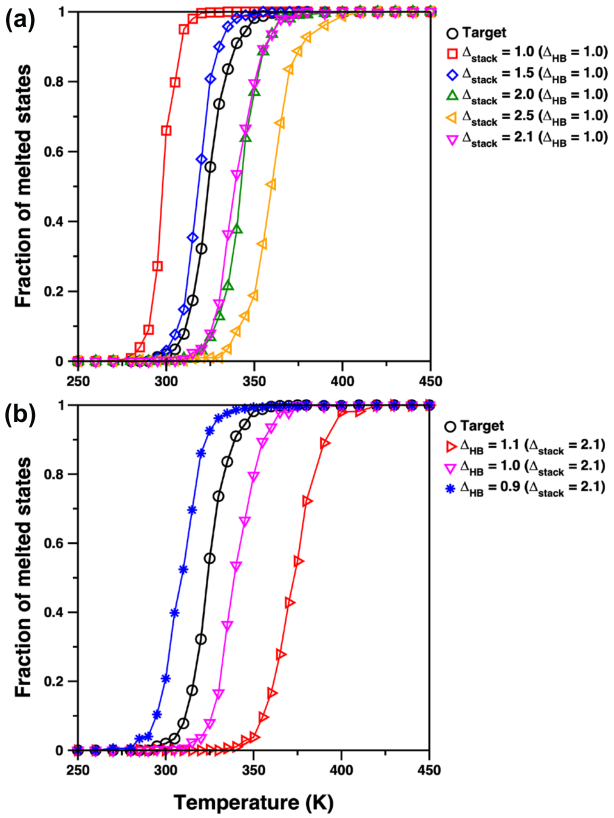


***Figure S1.*** Thermodynamic melting behavior for a pair of dsDNA S1S2: 5′-GCGTCATACAGTGC-3′ obtained from parallel-tempering simulations using our 2-bead CG DNA model, as the energy parameters are scaled for (a) stacking interactions, and (b) hydrogen bonding interactions (along with optimized stacking energy parameters).

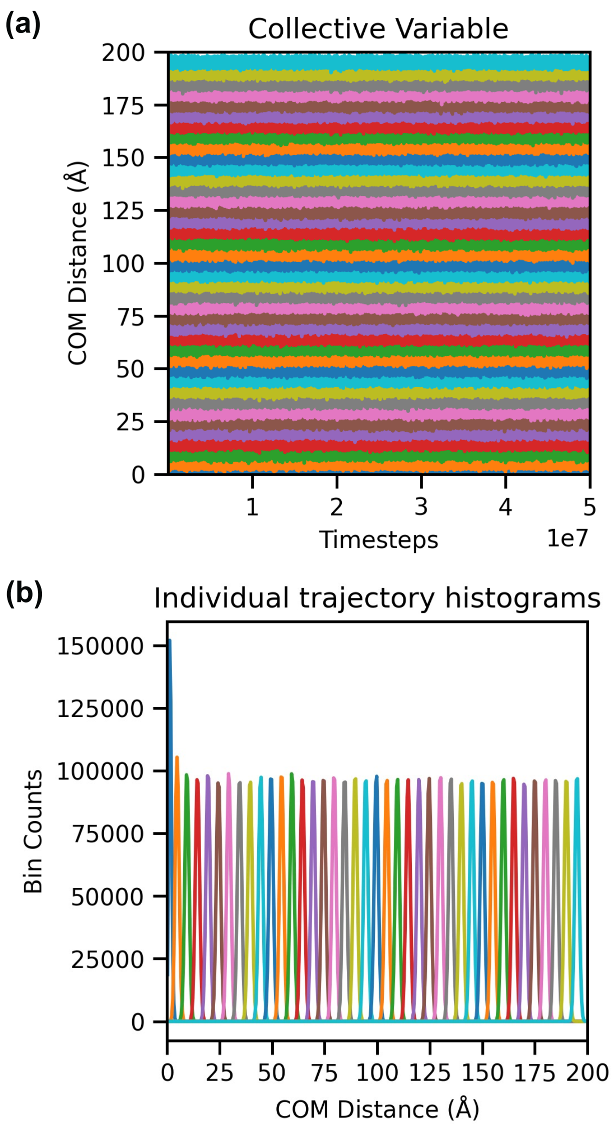


***Figure S2.*** (a) Collective variable as a function of simulation time, and (b) distribution of the regions that the collective variable samples (histogram per replica).

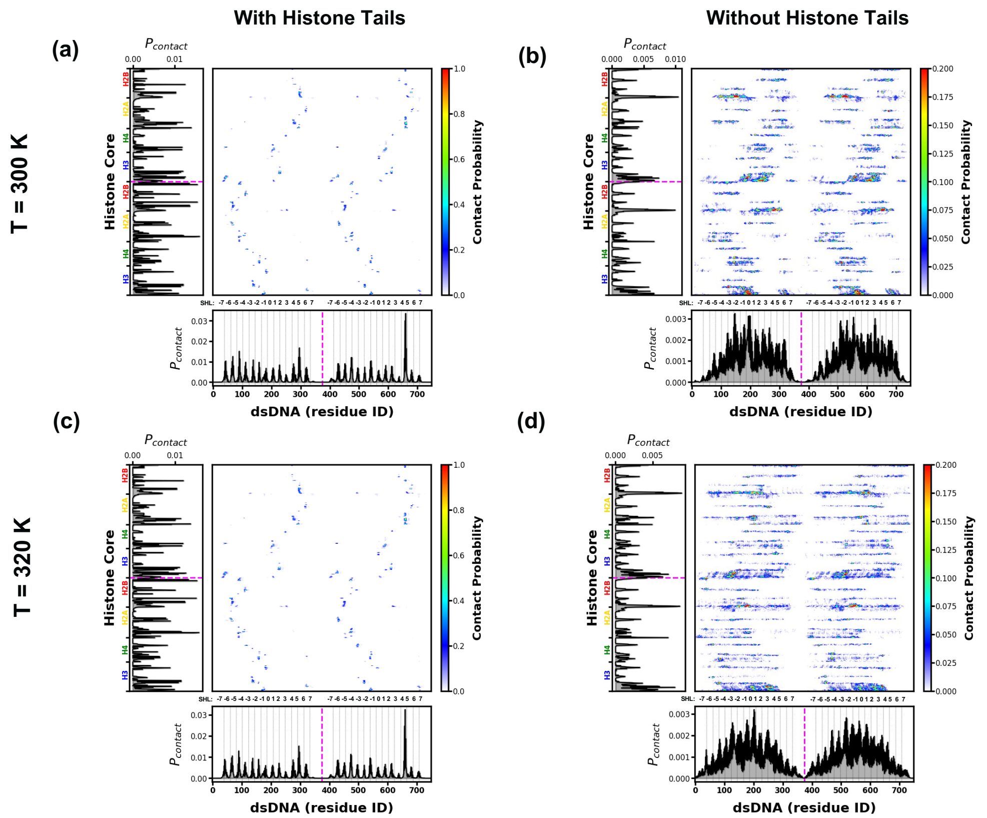


***Figure S3*.** Intermolecular contacts between DNA and histone core, for nucleosome with histone tails or without histone tails, respectively, when the nucleosome is **simulated at two different temperatures (T = 300 K versus T = 320 K)**. Preferential interactions are shown in red color. We note that the dotted magenta color lines on the x-axis and y-axis indicate the demarcation between two DNA strands or copies of histone core, whereas the gray color grid lines on the x-axis show superhelical locations (SHL).

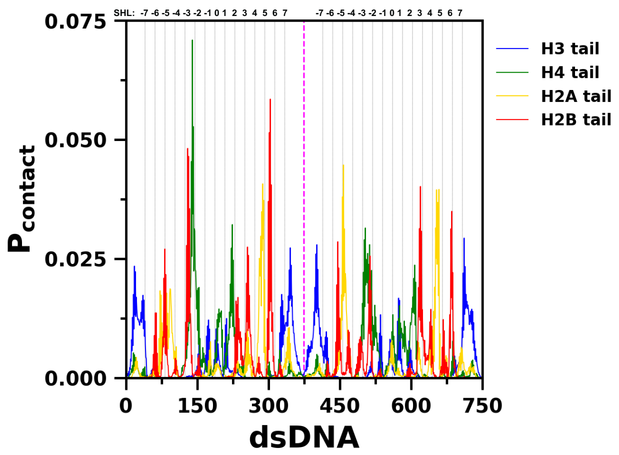


***Figure S4.*** The probability of intermolecular contacts between tails of histones and DNA. We note that the dotted magenta color line indicates the demarcation between two DNA strands and the grey color grid lines on the x-axis show superhelical locations (SHL).

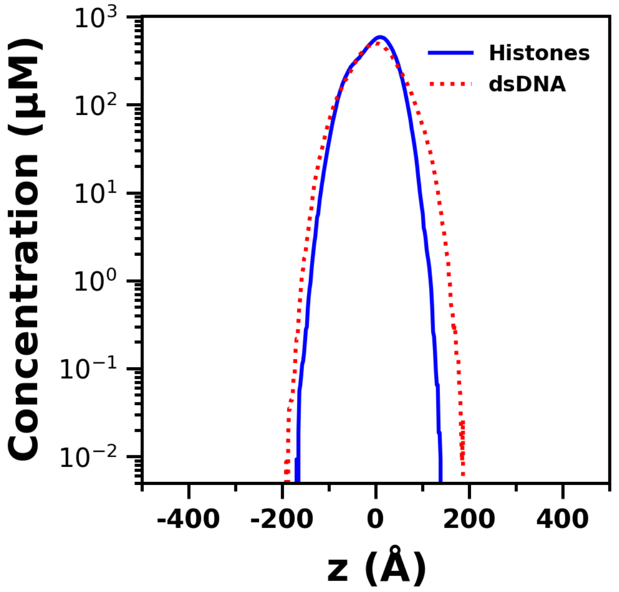


***Figure S5.*** Comparison of the concentration profiles of the histone proteins and nucleosomal DNA when the nucleosome is simulated with HP1α.

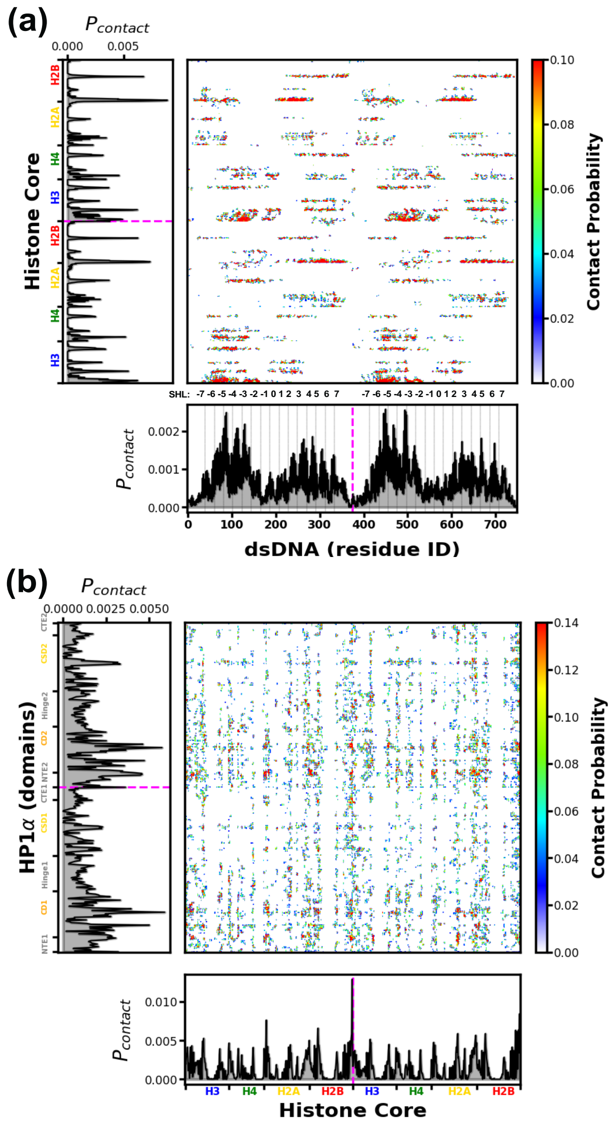


***Figure S6.*** Intermolecular contacts between (a) DNA and histone core, and (b) HP1α and histone core, **when the nucleosome is partitioned in HP1α droplet**. Preferential interactions are shown in red color. We note that the dotted magenta color lines on the x-axis and y-axis indicate the demarcation between two DNA strands or copies of histone core, whereas the gray color grid lines on the x-axis show superhelical locations (SHL).

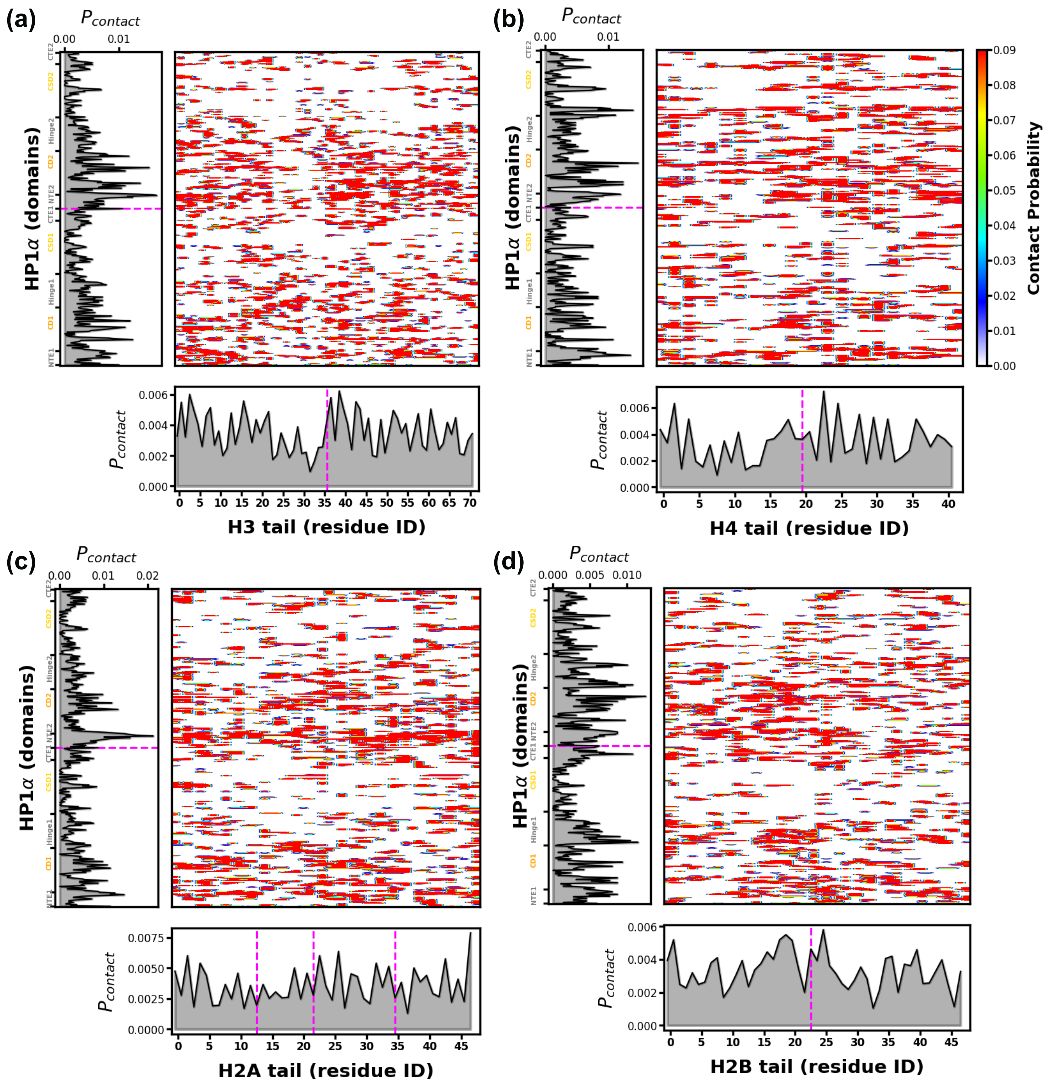


***Figure S7.*** Intermolecular contacts between HP1α and histone tails: (a) H3, (b) H4, (c) H2A, (d) H2B. Red color indicates preferential interactions. Dotted magenta color lines on the x-axis and y-axis indicate the boundaries between HP1α dimers, DNA strands, and histone tails.

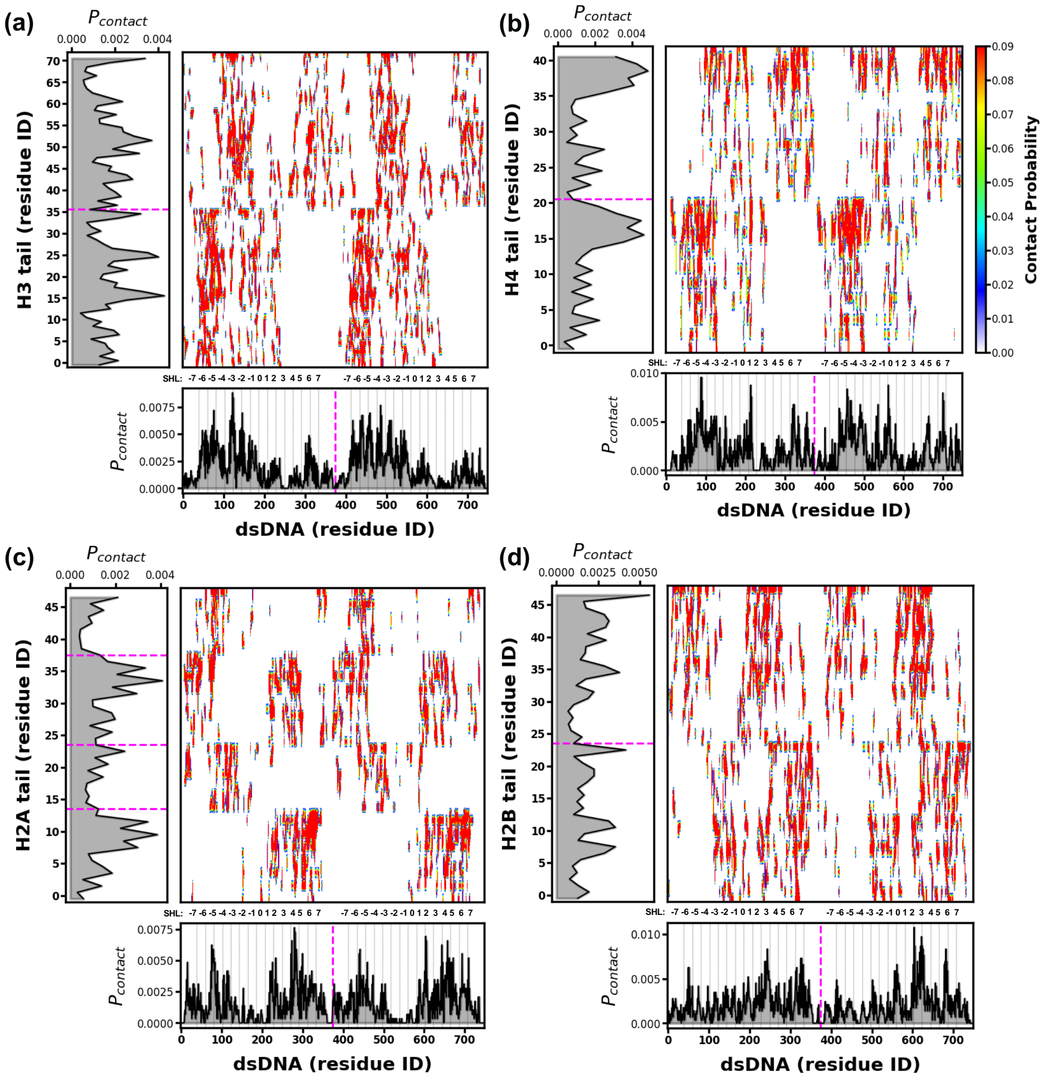


***Figure S8.*** Intermolecular contacts between DNA and tails of histones (a) H3, (b) H4, (c) H2A, (d) H2B, **when the nucleosome is partitioned in HP1α droplet**. Preferential interactions are shown in red. We note that the dotted magenta lines on the x-axis and y-axis indicate the demarcation between two DNA strands and the histone tails, respectively, whereas the grey color grid lines on the x-axis show superhelical locations (SHL). Please refer to **Figure 3b** of the main manuscript for SHL coordinate system considered in this work.
